# Extracellular biosynthesis of silver nanoparticles using cell suspension of *Rhodococcus fascians*

**DOI:** 10.1101/2024.03.27.587053

**Authors:** Farideh Ghadamgahi, Naga Charan Konakalla, Masome Mehraban Sang Atash, Rodomiro Ortiz, Ramesh Vetukuri

## Abstract

Biosynthesis of metallic nanoparticles using biological systems such as bacteria has become an important nanobiotechnology area. In this report, we present the first extracellular biosynthesis of silver nanoparticles (AgNPs) using the gram-positive bacterium *Rhodococcus fascians*. The AgNPs underwent characterization through various analytical techniques, encompassing UV–visible spectroscopy, X-ray diffraction (XRD), Fourier transform infrared spectroscopy (FTIR), Transmission electron microscopy (TEM) and Scanning electron microscopy (SEM). UV-visible spectroscopy revealed the emergence of an absorbance peak at 430 nm due to the synthesis of AgNPs. *R. fascians* started producing AgNPs after 12 h of incubation, with the highest yield after 48 h. The extent of synthesis was higher when cultures were grown in the dark than in the light. According to TEM and SEM, the AgNPs had a uniform spherical morphology of diameter 10–80 nm. The AgNPs exhibited antifungal efficacy against the virulent filamentous fungi *Rhizoctonia solani, Botrytis cinerea*, and *Fusarium graminearum*, which cause root rot, soft rot and head blight on plants, respectively. This research provides evidence on the ability of *R. fascians* to generate AgNPs from silver nitrate, as well as their subsequent assembly and potential for controlling vascular wilt disease.

## 1. Introduction

Over the past decade, there has been significant interest in nanoparticles (NPs) and nanomaterials due to their distinctive size-dependent physical and chemical properties [1,2]. Nanomaterials possess significant physical properties, among other characteristics, their wide surface area-to-volume ratio, and elevated surface energy. The synthesis of nanostructured materials and metal NPs holds appeal owing to their extraordinary photo-electrochemical, optical, electronic and chemical characteristics [3,4]. Silver nanoparticles (AgNPs), utilized in 383 products globally with an average annual production of 55 tons, stand as one of the most prevalent engineered nanomaterials [5]. Their primary application stems from their potent antimicrobial properties, effectively combating pathogens like fungi, bacteria and viruses [6,7]. Consequently, they find extensive use across various products, including detergents, medicine, food additives and many more[8,9]. The synthesis of silver nanoparticles is achievable through chemical, physical, and biological means [10,11]. Physical methods were initially utilized to synthesize metallic nanoparticles, whereas chemical methods now predominate [12]. However, nanoparticles produced via chemical methods entail the utilization of toxic chemicals such as sodium, sodium nitrate, and alcohol, serving as reducing and stabilizing agents [13]. Not only are these methods potentially hazardous due to their flammability, but they also yield non-degradable nanoparticles, contributing to environmental pollution. In light of these challenges and environmental concerns associated with silver nanoparticle production, the imperative arises for developing environmentally compatible, cost-effective, and chemical-free biological methods [14–17]. The biosynthesis of silver nanoparticles utilizing plant extracts and microorganisms such as, algae, bacteria, fungi and yeasts emerges as a promising alternative approach in this regard [18–21]. Such microorganisms are potential candidates for various applications because of their low toxicity, eco-friendly, high foaming property, high selectivity, and specific activity under extreme pH, temperature and salinity conditions [19,22].

Among different sources of nano particles, bacteria hold more interest due to their lower cost, rapid procedure and faster reproduction rates than fungi or plants [23,24]. Bacteria offer numerious examples for biosynthesis of AgNPs, such as *Pseudomonas alloputida* [25], *Bacillus cereus* [26], *B. pumilus* [27], *B. subtillus* and *B. megaterium* [28], *Pseudomonas spp* [29], and *Terrabacter humi* [30], and *Oscillatoria limnetica* [31]. In this study, we developed a biological method for synthesizing AgNPs using *Rhodococcus fascians*. Among the various applications, the antimicrobial effect [32] is most important focus of our study. *R. fascians* is a gram-positive, nonmotile, aerobic bacterium belonging to the actinomycetes group [33]. In this study, we focused on the biological approach for synthesizing AgNPs using *R. fascians*, under two distinctive conditions: dark and light. We evaluated their biocontrol efficacy against various filamentous fungal phytopathogens.

## 2. Materials and Methods

### 2.1. Materials

Silver nitrate (AgNO_3_) was purchased from Sigma Company (St. Louis, MO, USA). *R. fascians* bacteria and filamentous fungi *Rhizoctonia solani, Botrytis cinerea* and *Fusarium graminearum* (causative agents of root rot, soft rot and head blight, respectively) were obtained from the Swedish Agricultural University, Alnarp, Sweden.

### 2.2. Biosynthesis of AgNPs

For the biosynthesis of AgNPs, a single colony of bacterial *R. fascians* was initially cultured for 16 h in 300 ml of Muller Hinton broth in a 1000 ml Erlenmeyer flask at 28°C with agitation using a shaker at 150 rpm [31]. Afterwards, the culture was centrifuged at 10000 × g for 10 min, the supernatant was removed, and the cell pellet was washed three times with 1x phosphate-buffered saline (PBS). After collecting the biomass, 400 ml sterile water was mixed with 4 g of the bacterial pellet in a 1000 ml Erlenmeyer flask and incubated for 24 h at 28°C and 150 rpm. Subsequently, the suspension was centrifuged, and the pellet was discarded. Then, 40 ml of 1mM of AgNO_3_ solution was added to the mixture, which was evenly divided equally into two tubes for each designated incubation time: one was kept in the dark (covered with aluminium foil), and the other was kept in the light. The tubes were incubated for various incubation times up to 48 h at 28°C in an orbital shaker set to 150 rpm. A control was also prepared the same way but without AgNO_3_. The experiment was repeated three times. Various analytical techniques were employed to characterize the formation of AgNPs, including UV–visible spectroscopy, transmission electron microscopy (TEM), scanning electron microscopy (SEM), Fourier transform infrared spectroscopy (FTIR), and X-ray diffraction.

### 2.3. Characterization of Synthesized AgNPs

After incubating at various time points, 1 ml of the supernatants were prepared under dark, light, and control conditions and used for UV-Vis analysis. The formation of AgNPs was followed by recording UV-Vis spectra between the wavelengths 300 and 700 nm using a UV–visible spectrophotometer after 6, 12, 36, and 48 h. The biosynthesis of AgNPs was indicated by a change in colour of the solutions from initial pale yellow to reddish and the presence of an absorbance peak at 430 nm [34].

### 2.4. TEM Measurements

The size, shape, morphology, distribution and localization of AgNPs produced by *R. fascians* were determined by TEM. To prepare synthesized AgNPs for imaging by TEM, a droplet of suspension containing AgNPs was applied onto a carbonized copper grid and subsequently desiccated in a vacuum desiccator and captured using a transmission electron microscope (Zeiss, Germany) [15].

### 2.5. SEM and Energy Dispersive X-Ray Spectroscopy

SEM analysis was conducted at the Amirkabir University of Tehran (XL30 Philips WDX-3pc, Microspec cor., USA). To determine the atomic composition of the solutions, aluminium foil was mounted onto the support bases of the samples and cleaned with alcohol. A thin layer of gold was then sputter-coated onto the aluminium to prevent the formation of static charge during the analysis [15]. The SEM instrument was equipped with an energy-dispersive X-ray spectroscopy (EDX) system. EDX is a method of analysis used to characterize a sample’s chemical or elemental composition. It relies on interaction of an X-ray excitation source and a sample.

### 2.6. X-ray Diffraction

XRD is a technique utilized to identify the internal structure of crystals by analyzing the powder diffraction pattern of a target material. The multitude of crystals within the powder sample cause different crystal planes to be randomly exposed to radiation at different angles, with the diffraction intensity subsequently measured. This process enables the characterization of crystal structure and phase composition within the material. We employed XRD to determine the crystal structure and average size of the synthesized AgNPs. The samples used for TEM analysis were also subjected to XRD analysis. The prepared AgNPs solution was freeze dried at -46°C for 10 h, and the resultant powder was analyzed with an X-ray diffractometer (Unisantis XMD300, Germany). The powder XRD pattern was documented for a range of 2ϴ from 30° to 80° [36,37].

### 2.7. Fourier transform infrared (FTIR) spectroscopy

FTIR analysis was performed at room temperature using a Thermo Nicolet Avatar 370 spectrometer (Germany) to identify the functional groups and surface chemistry of dried AgNO_3_ and synthesized AgNPs in the range 400–4000 cm^−1^ with a resolution of 4 cm^−1^ [38,39].

### 2.8. Antagonistic Activity of Synthesized Nanoparticles

We have employed disk diffusion method to investigate the inhibitory effect of synthesized AgNPs which was previously reported on pathogenic fungi *R. solani, B. cinerea* and *F. graminearum* [40]. A 0.5 cm diameter sterile filter disk was drenched in 2 ml of AgNPs solution for 15 min and then placed in the edge of a Petri dish containing potato dextrose agar (PDA) medium (Merck, Darmstadt, Germany). A control dish comprising a disc drenched for 15 min in AgNO_3_ solution (1 mM) on PDA medium was also prepared. Subsequently, a 0.5 cm fungal plug from a fresh subculture was placed onto the center of plates and incubated at 25 ± 2°C for 7 days until the leading edge of the fungus in the control nearly reached the edge of the Petri dish. The antagonistic effect was quantified by measuring fungal growth and calculating the percentage inhibition rate using Eq. 1 [41]: Growth Inhibition % = (GC−GT)/GC×100. GT is the distance of fungal growth from fungal plug to the ledge of growth in the treatment samples and GC is the fungal growth in control plates. The experiment was performed using a completely randomized design with at least three replicates per pathogen. These experiments were-conducted three times.

### 2.9. Data analysis

All statistical data analysis were carried out using JMP Pro 17 statistical software. The antifungal activity of AgNPs were analysed by the analysis of variance (ANOVA) and compared by Student’t Test with twelve replicates.

## 3. Results

### 3.1. Extracellular Biosynthesis of AgNPs

To evaluate the extracellular synthesis of AgNPs by *R. fascians*, the supernatant of the isolate was mixed with AgNO_3_ solution under two different conditions: light and dark (Fig. 1). After 12 h shaking, the color changed to reddish, indicating the reduction of Ag+ ions to Ag° (AgNPs) (Fig. 2C and D). The final mixture under dark conditions had a stronger color than under light conditions (Fig. 2D).

**Fig 1.**
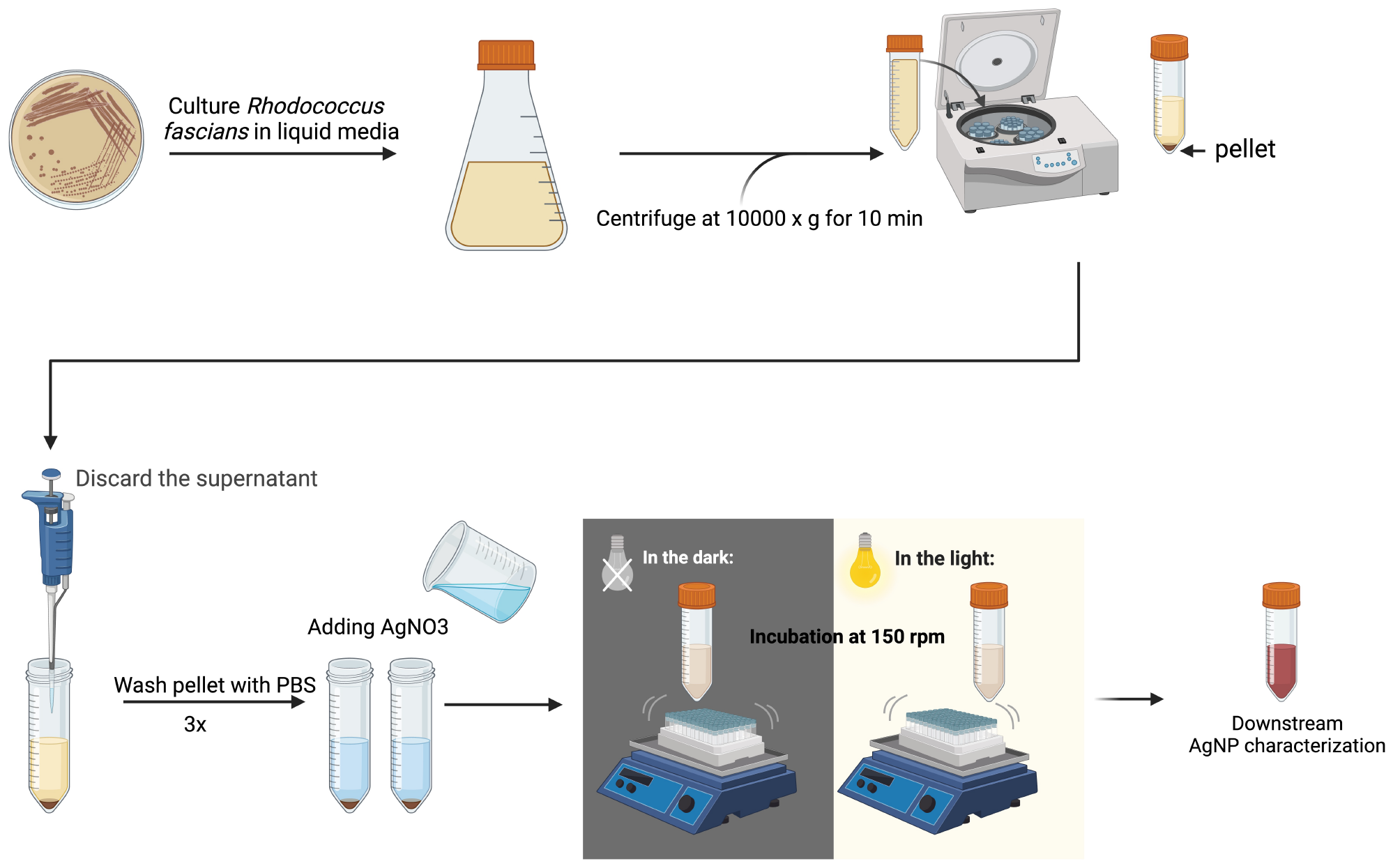
Steps involved in the biosynthesis of silver nanoparticles (AgNPs).

**Fig 2.**
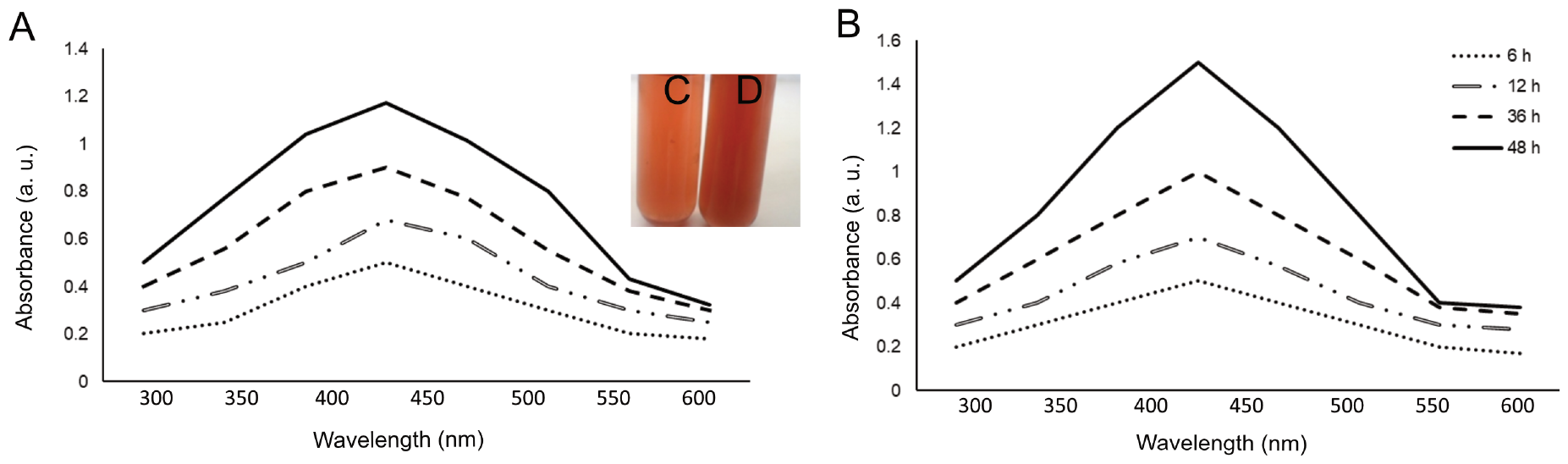
UV–visible absorption spectra of AgNPs synthesized by *Rhodococcus fascians* for different incubation times (6, 12, 36 and 48 h) under (A) light and (B) dark conditions. The inset in (A) shows a photograph of the AgNP solution after 48 h incubation under (C) light and (D) dark conditions.

### 3.2. UV–Visible Spectroscopy of Synthetic AgNPs

AgNPs solutions obtained after different incubation times of 6, 12, 36 and 48 h with AgNO_3_, and 1 ml of the supernatants was used for UV–Vis analysis between the wavelengths 300–700 nm. The UV–Vis spectra exhibited a prominent peak at 430 nm, indicating the presence of AgNPs [42]. The intensity of the absorbance band increased with longer incubation time, reaching its peak after 48 hours of incubation under both dark and light conditions (Fig. 2). However, only the solutions obtained in the dark were selected for further research due to their absorption peak was higher and sharper than under light conditions (Fig. 2). Previous studies have shown that a sharper peak indicates smaller NPs, whereas a higher absorption peak indicates a higher synthesis rate of NPs [43].

### 3.3. Characterization of AgNPs by TEM

TEM was employed to determine the size, shape, and spatial distribution of the particles [44,45]. Briefly, a drop of the sample diluted in distilled water was placed onto a carbonized copper grid and dried in a vacuum desiccator before observation by TEM. TEM images of AgNPs at four different scales (20, 50, 100 and 150 nm) are shown in Fig. 3A. The TEM analysis of the bacterial synthesized AgNPs indicated the creation of spherical nanoparticles ranging in size from 10–80 nm (Fig. 3). A particle size distribution histogram determined by TEM is shown in Fig. 2B. The majority of particles had diameters of 30–40 nm.

**Fig 3.**
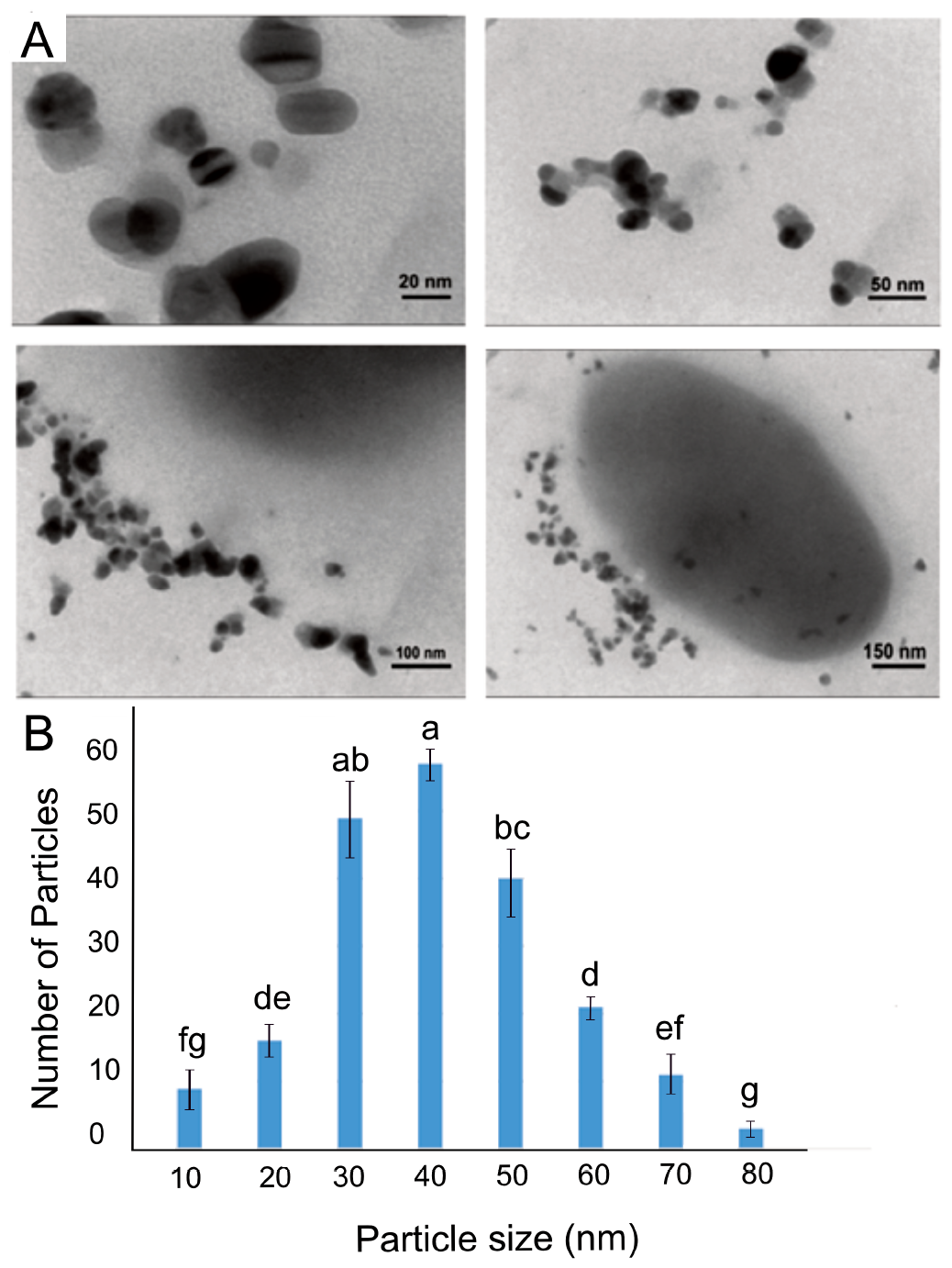
(A) Transmission electron microscopy images of AgNPs in *R. fascians* culture supernatant after 48 h incubation recorded at different scales (20, 50, 100 and 150 nm), (B) Histogram of the particle size distribution of AgNPs. Means labelled with different letters are significantly different according to Student’s t test at p < 0.01.

### 3.4. Characterization of AgNPs by SEM

SEM analysis showed that the synthesized AgNPs were dispersed with roughly spherical structures (Fig. 4A and B). The synthesized AgNPs were easily recognizable by SEM. ImageJ software was used to assess the diameters of 40 randomly selected particles, suggesting a size range of 10 to 70 nm. To analyze the elemental composition of the AgNPs, EDX was employed, which confirmed that the particles contained a high percentage Ag (Fig. 4C).

**Fig 4.**
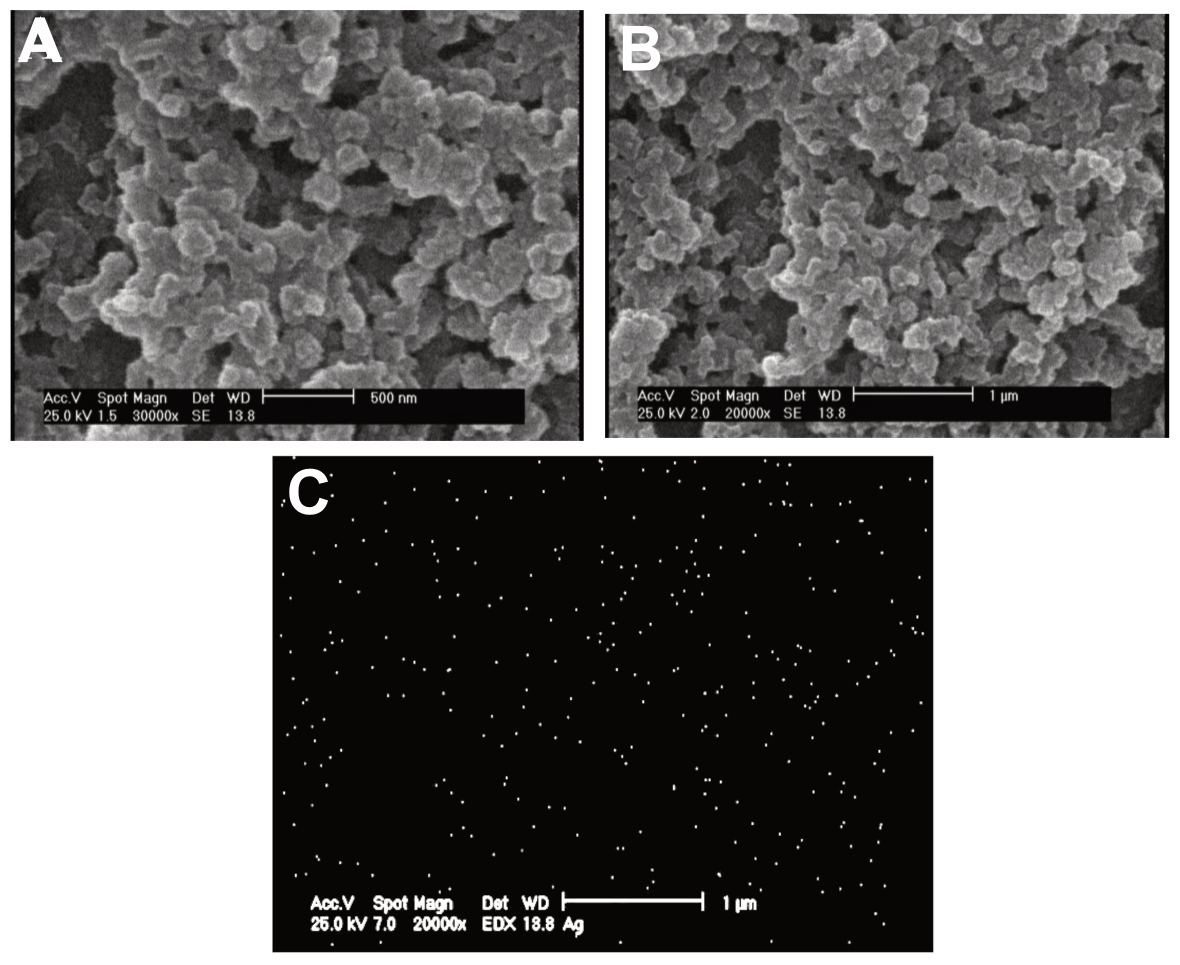
Scanning electron microscopy images of the self-assembly of AgNPs produced by *R. fascians* (A) 500 nm scale bar and (B) 1 μm scale bar. (C) Energy dispersive X-ray image representing Ag particles as white dots.

### 3.5. FTIR Analysis

FTIR measurements were performed to investigate putative interactions between the AgNPs and capping proteins, which may have been responsible for the decrease in Ag^+^ ions and subsequent stabilization of AgNPs (Fig. 5A and B). Amide bonds between amino acid residues in proteins produce a typical signature in infrared spectra. Fig. 5B shows an FTIR spectrum of the control (bacterial supernatant without AgNO_3_) with the following peak assignments: 3496 cm^-1^ (OH), 2091 cm^-1^ (CN) and 1642 cm^-1^ (carbonyl group). However, a new peak at 3265 cm^-1^ was observed in the spectrum of AgNPs synthesized by *R. fascians* due to the presence of an NH group (amino group) [46,47]. The emergence of absorption peaks in the range 700-1400 cm^-1^ may indicate the complexation of amide groups with AgNPs (Fig. 5A). The new small peaks at 3300 to 3500 cm^-1^ was likely due to stretching of OH groups in carbohydrates and H_2_O [37].

**Fig 5.**
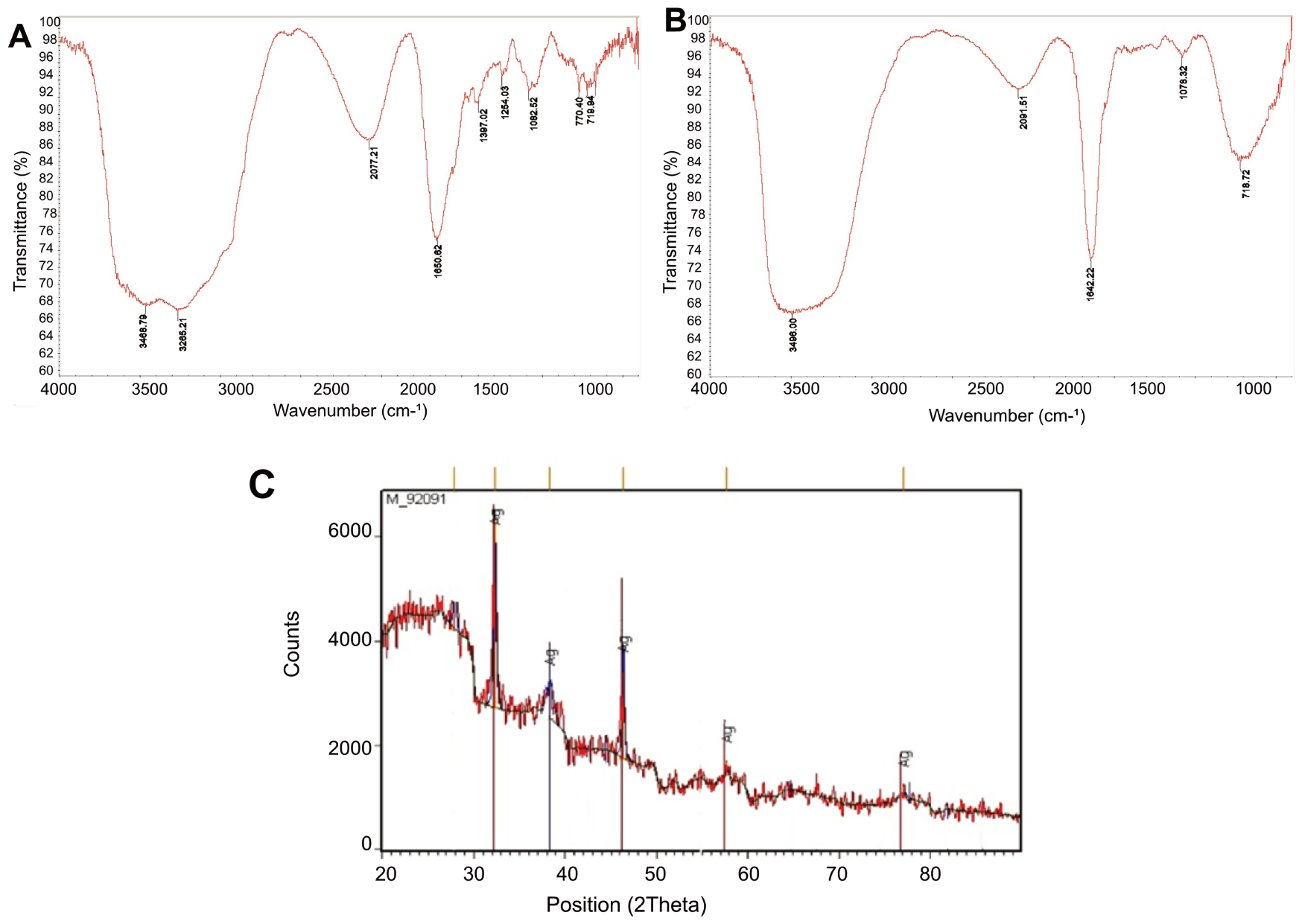
Fourier transform infrared spectroscopy absorption spectra of *R. fascians* (A) with AgNO_3_ (synthesized AgNPs), (B) without AgNO_3_. Differences are indicated in the peaks and assigned positions. (C) X-ray diffraction pattern of AgNPs synthesized by *R. fascians*.

The overall FTIR pattern confirmed that capping proteins were present in the AgNPs and that these proteins were not substantially aggregated. Interactions between proteins (e.g., enzymes) and NPs can be mediated by free amine groups, cysteine residues or electrostatic attraction of negatively charged carboxylate groups. Hence, the synthesis and stabilization of AgNPs maybe due to the free amine and carbonyl groups of bacterial proteins [48].

### 3.6. Characterization of AgNPs by XRD

Silver nanoparticles were synthesized (see materials and methods), were analyzed by XRD demonstrating the formation of AgNPs (Fig. 5C). The XRD analysis revealed the presence of four prominent peaks at specific 2ϴ values of 32.1°, 38.2, 46.2°, 57.4°, and 76.8°, regarding to silver crystals with 111, 200, 220, 311 and 331 structures, respectively. Other observed peaks may be attributed to crystallization of biological compounds present during the synthesis process, leading to the bio-mineralization of the NP surface. The XRD analysis confirmed that AgNPs were formed by *R. fascians*, and the sharpness of the peaks indicated that the particles had a crystalline structure [49].

### 3.7. Antifungal Effects of AgNPs

Fungi *R. solani, B. cinerea* and *F. graminearum* were treated with synthesized AgNPs using the disc diffusion method on PDA plates (Fig. 6). The fungal growth inhibiting rate was determined from Eq. 1 (see materials and methods) by measuring the diameter from the center of the Petri dish to the leading edge of the fungus. *F. graminearum* showed the highest percentage inhibition (41.66%), followed by *R. solani* (29.63%) and then *B. cinerea* (15.92%). The results demonstrated therefore that growth of *R. solani, B. cinerea* and *F. graminearum* was inhibited in the presence of AgNPs produced by *R. fascians*.

**Fig 6.**
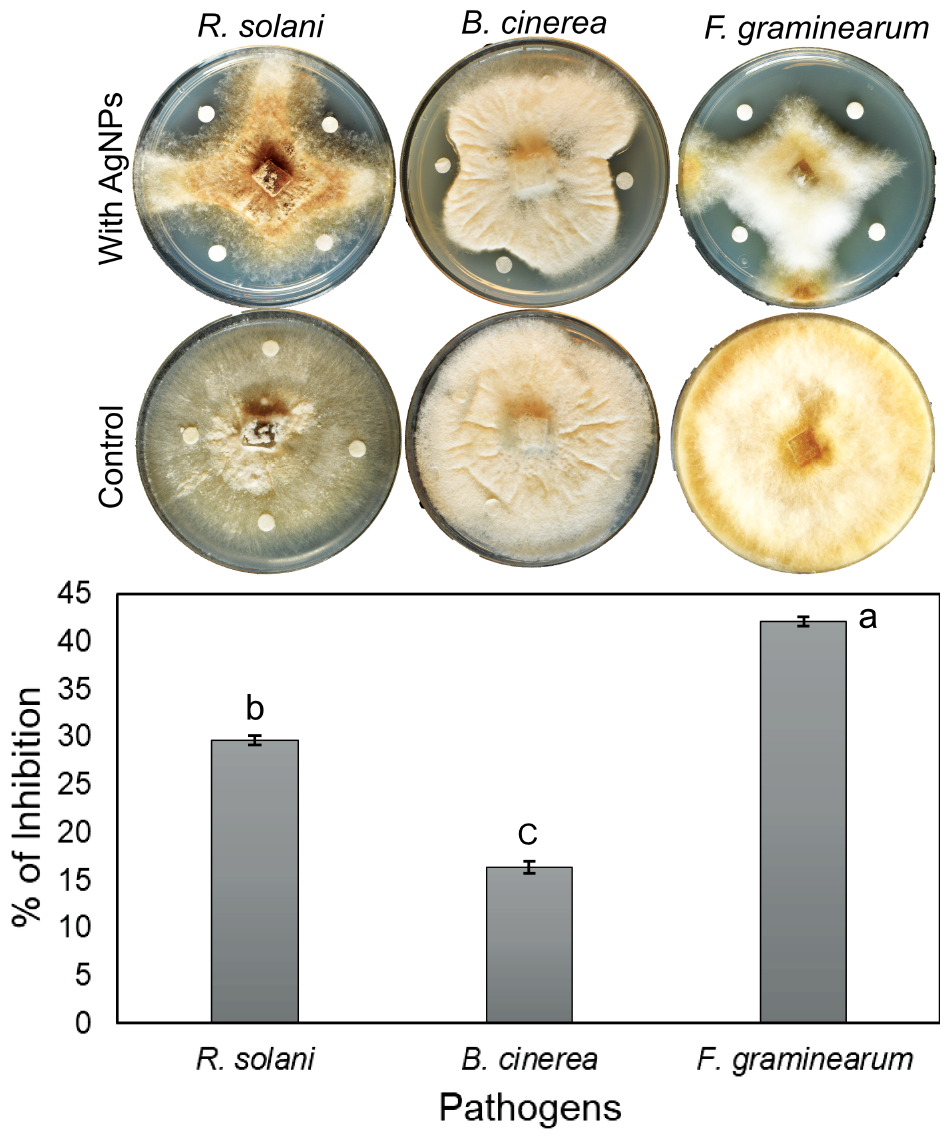
Antifungal activity of synthesized AgNPs against *R. solani, B. cinerea and F. graminearum*. Data represent means of 12 replicates for each treatment. Means labelled with different letters are significantly different according to Student’s t test at p < 0.01.

## 3. Discussion

Biosynthesizing AgNPs is more affordable and environmentally friendly than chemical and physical procedures. During biosynthesis, bacterial cells and metabolites serve as stabilizing and reducing agents, enabling the facile reduction of silver ions to produce stable AgNPs with well-defined size and shape. The extracellular formation of AgNPs could offer advantages in industrial preparations by minimizing the number of steps required for their purification. So far, bacteria such as *B. cereus* [50], *B. licheniformis* [14], and *Streptomyces sp*. ERI-3 [51] generated extracellular AgNPs. The current investigation contributes to this list by identifying *R. fascians*, a novel producer of AgNPs, a species not previously reported to be able to produce AgNPs. It has been proposed that AgNPs are synthesized by enzymatic reduction mediated by NADPH-dependent nitrate reductase in the microorganisms. This enzyme has been shown to convert silver ions to metallic silver when nitrate ions activate it [52]. Following the colour shift, it has been hypothesized that DNA, sulfur-containing proteins and NADH-dependent nitrate reductase are involved in producing AgNPs by bio-reducting silver ions to metallic silver. Previously, *B. licheniformis* and *B. cereus* have been used to reduce silver ions in a biomimetic method for producing Ag nanocrystals [14,31,52].

The present study reported that *R. fascians*, could induce the synthesis of AgNPs in the culture supernatant. Therefore, it consists of components necessary for producing AgNPs that must exist inside *R. fascians* cells. UV-Vis spectroscopic analysis confirmed the conversion of silver ions to AgNPs according to the presence of an absorption peak at 430 nm as well as a colour change (Fig. 2). Previous research demonstrated that spherical Ag-NPs contribute to absorption bands in the UV-visible spectrum at about 400-450 nm [44]. In this study, AgNPs were synthesized under either light or dark conditions. According to preliminary results, UV-Vis absorption peaks at 430 nm and color changes were more prominent in solutions kept in the dark than in the light. However, the mechanism by which darkness influences the formation of AgNPs is not yet known (Fig. 2).

SEM analysis of the *R. fascians* produced AgNPs showed that they were spherical with an average particle size of 10 to 70 nm (Fig. 4). In the present study, TEM showed that the production of AgNPs was extracellular and confirmed the size and shape of particles observed by SEM (Fig. 3). FTIR spectroscopy indicated that the stability of the AgNPs was likely due to the presence of bacterial proteins and enzymes coating the particles. XRD and EDX results confirmed that the main element of the NPs was silver (Fig. 5). In the test of antifungal activity, the findings showed that AgNPs produced by *R. fascians* inhibited the growth of *R. solani, B. cinerea*, and *F. graminearum* (Fig. 6).

Previous research examined the antifungal activity of AgNPs against various plant pathogenic fungi [20,39,53]. For example, the fungicidal activity of AgNPs produced by *Trichoderma asperellum* was investigated against four soil-borne plant pathogens, *R. solani, F. oxysporum, Sclerotinia sclerotiorum* and *S. rolfsii*. The results revealed that AgNPs showed significantly high efficacy in inhibiting the mycelial growth of the pathogens [53]. Additional research showed that NPs produced by a plant extract (*Chenopodium album* L) caused a 92% retardation in the fungus *Aspergillus terreus* [54]. However, the use of bacteria may be a more suitable, cheaper and faster option than plant extracts because of their rapid growth and lack of requirement for a specific culture medium. Also, silver nanoparticles may interact with proteins and other components of microbial cell membranes, causing membrane damage. Increased cell absorption and DNA fragmentation result in cell death and microbial growth suppression [26]. Ibrahim et al. [26] employed the disc diffusion method. They reported that NPs synthesized by *B. cereus* exhibited antimicrobial effects on bacterial strains *Staphylococcus epidermidis, Staphylococcus aureus, E. coli, Salmonella enterica, Porteus mirabilis*.

The microbial activity of AgNPs affects the structure of the bacterial/fungal cell. Alternatively, AgNPs may be incorporated into the cell wall, altering the membrane permeability and affecting respiratory processes [47]. Additionally, NPs might enter further into the microbial/fungal cell. Small AgNPs may likely have a more significant fungicidal effect than larger particles because of their larger surface area to volume ratio. According to other findings, positively charged silver ions are electrostatically attracted to negatively charged microbial membranes [55] and may be responsible for the antibacterial properties of AgNPs [56]. This may enable AgNPs to be used as advanced anti-microbial agents, drug delivery systems and nano-medical applications, including silver-based coatings, silver-layered medical devices, nano-lotions and nano-gels.

## 4. Conclusion

This study showed that *R. fascians* can initiate the synthesis of AgNPs after 12 h of incubation, with the highest synthesis rate achieved after 48 h of incubation. A characteristic absorbance peak at 430 nm was observed at all incubation times. The AgNPs exhibited a uniform spherical morphology with diameters of 10–80 nm. This report presents evidence for the first time on the ability of genus *Rhodococcus* to convert silver nitrate to AgNPs as well as their subsequent self-assembly.

